# Tolerance for heat stress in *Lemna minor* reflects both local adaptation and acclimation over time

**DOI:** 10.64898/2026.08.23.746491

**Authors:** Mario Blanco-Sánchez, Sonia E. Sultan, Koen J.F. Verhoeven

## Abstract

Assessing intraspecific variation in thermal stress tolerance is key to predicting plant responses and long-term persistence under climate change, yet its underlying sources and temporal dynamics remain poorly understood. Using a common garden experiment with four ecologically-relevant temperatures, we evaluated the sources and temporal dynamics of variation in temperature stress tolerance of 18 *Lemna minor* clonal lines from contrasting climates. Our results showed that past adaptation, physiological acclimation, and within-line variation jointly contributed to variation in performance. The study provides the first evidence of adaptive genetic differentiation in heat stress tolerance in this ecologically-widespread freshwater species, with lines from warmer regions showing higher growth under heat stress. However, these differences were transient and diminished under prolonged exposure. Experimental lines also showed acclimation over time, but these responses were strongly temperature-dependent and occurred only under sub-optimal conditions. Additionally, replicates from some lines exhibited divergent performance trajectories under sustained heat stress, suggesting the emergence of novel phenotypic variation, potentially mediated by epigenetic mechanisms. These results show that heat stress tolerance in *L. minor* arises from multiple interacting sources and is dynamically shaped by both selective history and immediate exposure time, suggesting a more nuanced, multi-layer understanding of variation in heat stress tolerance.

## 1. Introduction

Climate change is one of the most serious threats to biodiversity worldwide (Lovejoy & Hannah, 2019; Sage, 2020). Global temperature and precipitation patterns are changing rapidly, with an increased frequency of extreme events including droughts and heatwaves and a predicted global temperature increase of 1°C to 5°C (depending on greenhouse gas emissions) by the end of the century (IPCC, 2023). Consequently, global warming is threatening not only populations adapted to colder temperatures, but also those that currently thrive under thermal conditions close to their physiological maxima (Somero, 2010; Sage, 2020). Understanding sources of adaptive intraspecific variation in heat stress tolerance is critical to evaluate the potential for plant populations to persist in the face of climate change.

One possible source of such variation arises from past evolution in response to local conditions (Westerband *et al*., 2021; Blanco-Sánchez *et al*., 2024; Ramos-Muñoz *et al*., 2024). Widely distributed species encounter environmental gradients across their range, and such contrasting conditions often impose differential selective pressures (Hoffmann & Sgró, 2011; Blanquart *et al*., 2013). In the presence of genetically based phenotypic variation, and when the magnitude of natural selection is stronger than the effects of neutral evolutionary processes, differential selection will lead to adaptive differentiation (e.g., Blanquart *et al*., 2013; Sork, 2016). As a result, past adaptation to local temperature conditions often leads to clinal variation in heat tolerance, with genotypes from heat-prone regions showing higher performance under heat stress (e.g., Hoffmann *et al*., 2002; Woudstra *et al*., 2024; Lush *et al*., 2025). Indeed, significant associations between local climatic conditions and genetically-based phenotypic differentiation (i.e., phenotypic differences measured under common conditions) provide robust signals of adaptation to such environmental variation (Ramírez-Valiente *et al*., 2022; Anderegg, 2023; Blanco-Sánchez *et al*., 2024).

A second potential source of adaptive variation in heat stress tolerance is environmentally induced phenotypic change or phenotypic plasticity, which can enhance overall performance across conditions without requiring genetic change (Nicotra *et al*., 2010; Sultan, 2021). Among plastic responses, acclimation comprises a temporally dynamic form of plasticity involving gradual physiological adjustment to sustained environmental conditions. Noting that the term has been used in different ways in the literature (Leroi *et al*., 1994; Lagerspetz, 2006; Malinski *et al*., 2024), here we use *acclimation* to refer to physiological adjustments that maintain or enhance performance over time within a given environment (IUPS Thermal Commission, 2001; Bowler, 2005; Loeschcke & Sørensen, 2005). Acclimation to elevated temperatures has been widely documented in plants and involves physiological adjustments such as changes in photosynthesis and respiration, stabilization of cellular membranes, and the activation of stress-response pathways, including phytohormone signaling (e.g., ABA and SA), induction of heat-shock proteins and other molecular chaperones, antioxidant and ROS-scavenging systems, and the accumulation of osmolytes (Wahid *et al*., 2007; Larkindale & Vierling, 2008; Kumarathunge *et al*., 2019; Seydel *et al*., 2025; Wu *et al*., 2025). Importantly, genotypes may differ in their speed and ability to acclimate to environmental changes, such that acclimation responses can themselves evolve through natural selection (Matesanz *et al*., 2010; Richards *et al*., 2017). As a result, environmentally driven phenotypic variation can both buffer populations against short-term environmental fluctuations and facilitate their long-term persistence under climate change (Chevin & Hoffmann, 2017; Fox *et al*., 2019). However, despite the recognized role of acclimation in thermal stress tolerance, little is known about genetic variation in natural systems for the effectiveness and timing of acclimation responses to stressful temperatures.

Evaluating intraspecific variation in heat stress tolerance is particularly critical in plants, which have limited ability to migrate to track suitable temperature conditions as global climates warm (Jump & Peñuelas, 2005; Franks *et al*., 2014; Wang *et al*., 2023). Clonal plants are ideal systems to investigate potential sources and temporal dynamics of such tolerance, since genetically identical individuals can be grown under contrasting temperatures for multiple (asexual) generations. Due to its unusually short doubling time, the common duckweed (*Lemna minor* L.) has become a model clonal species for physiology, evolutionary ecology, and ecotoxicology studies (Ziegler *et al*., 2015; Laird & Barks, 2018; Gillies *et al*., 2024). The wide thermal distribution of the species makes it particularly suitable to evaluate the extent to which variation in heat stress tolerance arises from past adaptation to climate, environmentally-induced phenotypic acclimation, or a complex combination of both.

To gain insight into the sources of variation for temperature stress tolerance in this widespread species, we investigated how the performance of different clonal lines of *L. minor* changed over time across a set of ecologically relevant temperatures. We used a large, geographically broad sample of *L. minor* clones to assess quantitative genetic variation in heat stress tolerance, subsequently testing whether among-clone genetic differences matched climatic variation in their sites of origin (indicating past adaptation to local climates). In a common garden experiment, we assessed performance of 18 clonal lines in 4 temperature treatments (18°, 23°, 28° and 33°C), chosen to represent natural temperature variation among warmer *L. minor* populations during the species’ most active growing period (late-spring and summer) and temperatures projected in the near future for such locations due to climate change (Woolway *et al*., 2020; IPCC, 2023). Specifically, we addressed the following questions: i) Is there genetic variation in thermal stress tolerance among clonal lines of *L. minor*? ii) If so, does this genetic variation reflect adaptation to local climatic conditions at the sites of origin? iii) Do *L. minor* plants exhibit acclimation responses, expressed as increased performance over time within a given temperature? and iv) Do clonal lines differ in their acclimation responses to specific temperature conditions? Addressing these questions allows us to assess the role of genetic differentiation and acclimation in shaping temperature stress tolerance in *L. minor*.

## 2. Materials and methods

### 2.1 Species description and clonal lines evaluated

*Lemna minor* L. (Araceae), the common duckweed, is a floating freshwater aquatic plant inhabiting static bodies of water such as ponds and lakes as well as slow-current streams and canals. The species has a broad, near-cosmopolitan distribution excluding Arctic and Antarctic regions (Landolt, 1986, 1992; Laird & Barks, 2018; Tippery & Les, 2020). Plants of *L. minor* very rarely flower under laboratory conditions, instead showing clonal reproduction (Laird & Barks, 2018). *Lemna minor* has a doubling time of ∼2-5 days depending on both environmental conditions and genetic variation in growth potential (Ziegler *et al*., 2015; Laird & Barks, 2018).

To evaluate the sources of temperature stress tolerance in *L. minor*, we established a common garden experiment in the facilities of the Netherlands Institute of Ecology (NIOO-KNAW) that included 18 different *L. minor* clonal lines. These lines spanned a broad geographic and climatic range within the species’ northern hemisphere distribution, with a difference of ∼32° in latitude and ∼11 °C in mean temperature of the warmest quarter (when plants are mostly active and most growth occurs) between the coldest and the warmest location (Fig. 1; Table 1). Experimental lines were obtained from three duckweed stock collections from research groups based on Belgium (SCK CEN), Germany (Friedrich Schiller University Jena), and Italy (IBBA CNR).

**Figure 1:**
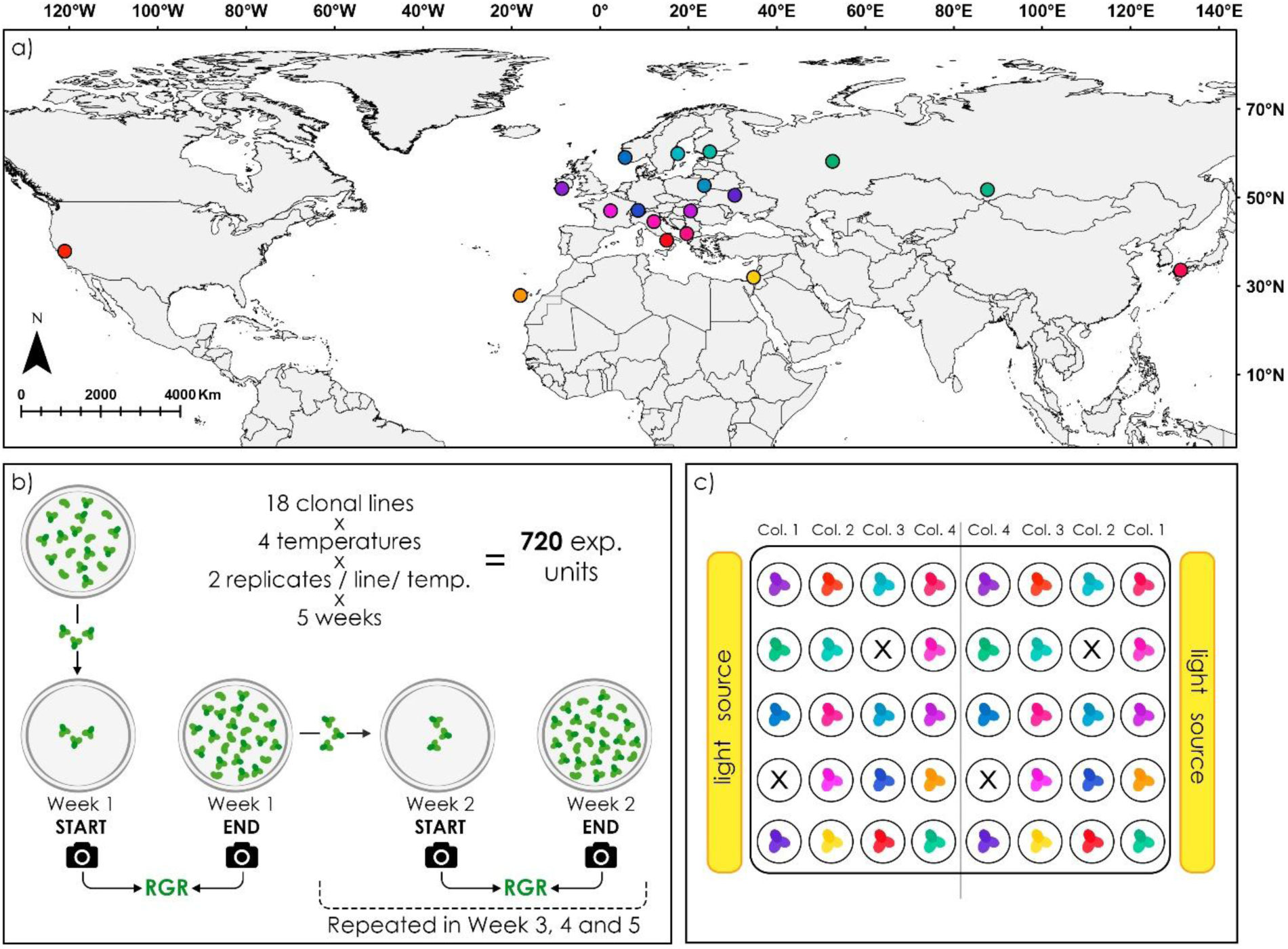
a) Source locations of the 18 clonal lines studied. Clonal lines are represented by different colors that reflect variation in temperature of origin, with slight adjustments to color palette to enhance visualization (i.e., warmer vs cooler colors denote lines from warmer vs cooler locations; see Table 1). This palette is maintained throughout all figures; b) Schematic representation of the experimental design. Eighteen *Lemna minor* clonal lines were grown under four constant temperature treatments (18 °C, 23 °C, 28 °C, and 33 °C), with two replicates per line and temperature, for five consecutive weeks (N = 720 experimental units). Each week, the relative growth rate (RGR) of each replicate was calculated from the change in total plant area over seven days; c) Spatial arrangement of jars within each growth chamber (see *Methods* for spatial blocking details). The first replicate of each line was randomly assigned to a 4 × 5 grid (block), and this arrangement was mirrored laterally in a second spatial block, creating an 8 × 5 grid such that each line had one replicate positioned near the chamber walls and one in the inner area, thereby balancing light exposure and any other spatial effects. Different clonal lines are represented with different colors, matching those in panel a).

**Table 1:**
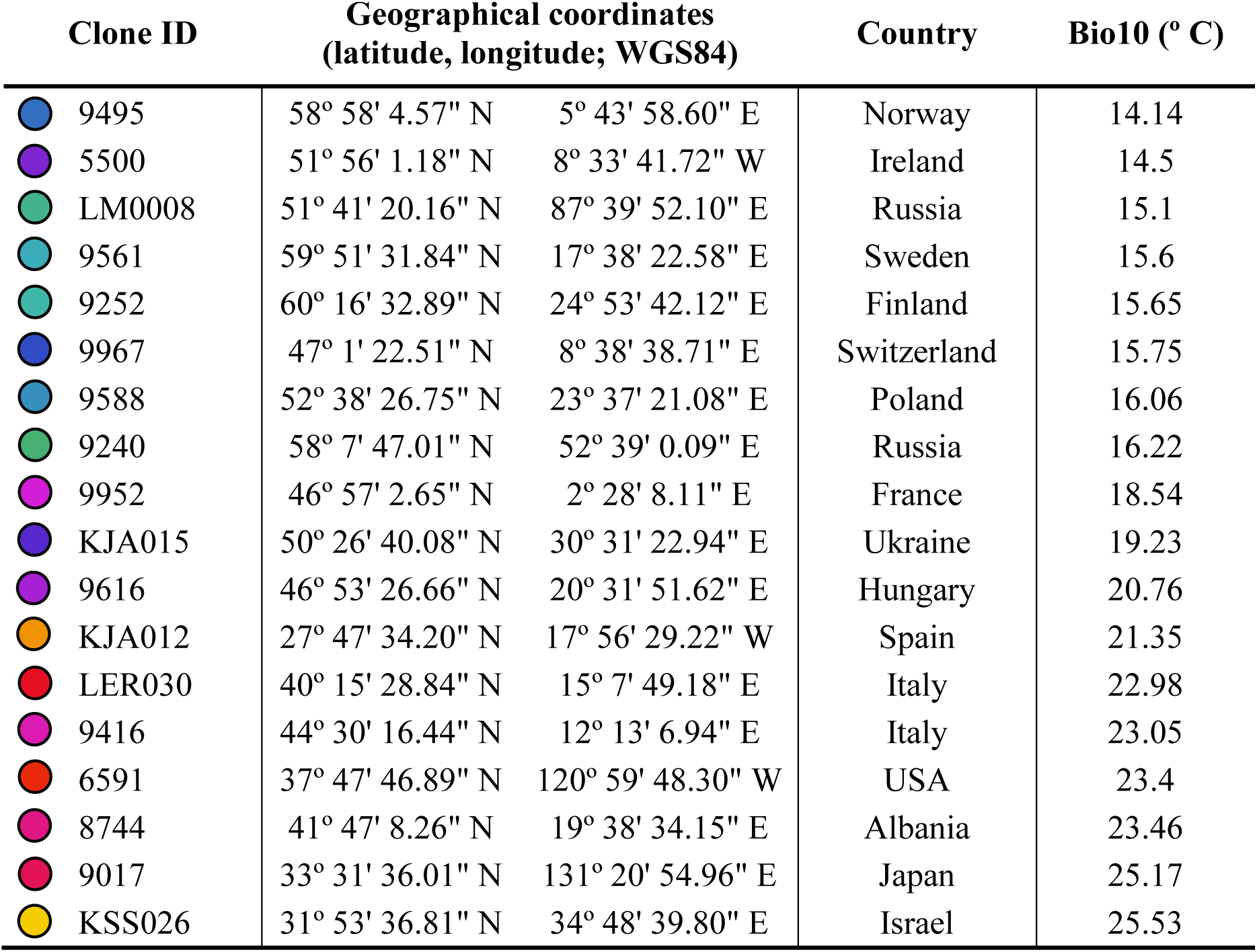
Clone IDs for the 18 experimental lines of *Lemna minor* along with their geographical coordinates, country of origin, and mean temperature of the warmest quarter (Bio10; climatic data extracted from WorldClim bioclimatic layers (Fick & Hijmans, 2017)). Colored circles correspond to the palette used throughout the figures, chosen to indicate colder vs warmer temperature of origin (ordered by Bio10).

| Clone ID | Geographical coordinates<br>(latitude, longitude; WGS84) |  | Country | Bio10 (° C) |
| --- | --- | --- | --- | --- |
| 9495 | 58° 58' 4.57" N | 5° 43' 58.60" E | Norway | 14.14 |
| 5500 | 51° 56' 1.18" N | 8° 33' 41.72" W | Ireland | 14.5 |
| LM0008 | 51° 41' 20.16" N | 87° 39' 52.10" E | Russia | 15.1 |
| 9561 | 59° 51' 31.84" N | 17° 38' 22.58" E | Sweden | 15.6 |
| 9252 | 60° 16' 32.89" N | 24° 53' 42.12" E | Finland | 15.65 |
| 9967 | 47° 1' 22.51" N | 8° 38' 38.71" E | Switzerland | 15.75 |
| 9588 | 52° 38' 26.75" N | 23° 37' 21.08" E | Poland | 16.06 |
| 9240 | 58° 7' 47.01" N | 52° 39' 0.09" E | Russia | 16.22 |
| 9952 | 46° 57' 2.65" N | 2° 28' 8.11" E | France | 18.54 |
| KJA015 | 50° 26' 40.08" N | 30° 31' 22.94" E | Ukraine | 19.23 |
| 9616 | 46° 53' 26.66" N | 20° 31' 51.62" E | Hungary | 20.76 |
| KJA012 | 27° 47' 34.20" N | 17° 56' 29.22" W | Spain | 21.35 |
| LER030 | 40° 15' 28.84" N | 15° 7' 49.18" E | Italy | 22.98 |
| 9416 | 44° 30' 16.44" N | 12° 13' 6.94" E | Italy | 23.05 |
| 6591 | 37° 47' 46.89" N | 120° 59' 48.30" W | USA | 23.4 |
| 8744 | 41° 47' 8.26" N | 19° 38' 34.15" E | Albania | 23.46 |
| 9017 | 33° 31' 36.01" N | 131° 20' 54.96" E | Japan | 25.17 |
| KSS026 | 31° 53' 36.81" N | 34° 48' 39.80" E | Israel | 25.53 |

### 2.2 Common garden experiment

Prior to the experiment, stock clonal lines were maintained in 60 × 15 mm Petri dishes containing 0.8% agarized SH medium (Schenk & Hildebrandt, 1972; prepared from Schenk & Hildebrandt Basal Salt Medium, Duchefa Biochemie, the Netherlands) and 0.1% of sucrose. Petri dishes were stored in a Snijders Micro Clima ECL02 growth chamber (Snijders Labs, the Netherlands) at 16 ± 0.2 °C and 50 ± 10 μmol m^−2^ s^−1^, under a light/dark photoperiod of 16/8 hours. Under such conditions, duckweed plants slow their growth, remaining viable for multiple months. Two weeks before the start of the experiment, three plants per clonal line were transferred into sterile glass jars with 150 mL of fully concentrated, sterilized N-medium (Appenroth et al., 1996). Clonal lines were kept separated and grown for two weeks at 22 ± 0.2 °C in another growth chamber of the same model, under 100 ± 10 μmol m^−2^ s^−1^ and a 16/8 h light/darkness photoperiod. This pre-experimental phase minimized potential storage effects and allowed the production of sufficient fully developed, healthy plants to set up the common-garden experiment.

On June 14^th^, 2024, three plants (each consisting of three fronds) from each clonal line were transferred to each of eight sterilized glass jars containing 150 mL of full-strength, sterilized N-medium (i.e., 24 plants per clonal line distributed in eight jars; eight jars per line x 18 clonal lines = 144 jars in total). To avoid algal contamination, experimental jars were covered with sterilized 60 × 15 mm Petri dishes that allowed minimal airflow between the inside and the outside of the jar. Immediately after transfer, plants in each jar were photographed using a Logitech Brio Ultra 4K webcam mounted in a fixed position at the top of a Godox LED Mini Photography Studio LST40 (Godox Photo Equipment Co., Shenzhen, China). The eight jars per clonal line were assigned to four Snijders growth chambers set at constant temperatures of 18, 23, 28, and 33 °C (± 0.2 °C), each with a 16/8 h light/dark photoperiod and 100 ± 10 μmol m⁻² s⁻¹ light intensity (two replicate jars per clonal line in each temperature). Within each growth chamber, jars were arranged in 2 mirror-image spatial blocks (1 plant per clonal line per block) on a single middle shelf (Fig. 1c). This experimental layout ensured an even spatial distribution and balanced exposure to light for experimental replicates within each treatment. Plants were grown for 7 days and photographed again. Then, three randomly selected plants from each experimental jar were transferred to new sterilized jars with 150 mL of fully concentrated, sterilized N-medium, which generated the two replicates per clonal line used in the following week. The experiment was run for five weeks, with weekly transfers and photos (*N* = 720; 18 clonal lines × 4 temperatures × 2 replicates/line/temperature × 5 weeks). All replicates of all clonal lines completed the experiment.

To characterize the performance of the experimental lines over time and across temperatures, we used image analysis to calculate the relative growth rate (RGR) of each replicate in each temperature and week based on the total surface area of the plants in each jar. Specifically, from the photos taken just after initial plant transfer and after one week of growth, the total area of the plants in each experimental jar was estimated using an AI instance model trained in Arivis Cloud (Zeiss Group, Germany). From these area measurements, we calculated RGR of each replicate in each temperature for each week as RGR = [ln(*A*_2_) − ln(*A*_1_)] / *T*_2-1_, where *A*_1_ is the area right after transferring, *A*_2_ the area after one week of growth, and *T*_2-1_ is the time elapsed between pictures (7 days).

### 2.3 Statistical analyses

All analyses were performed using R v4.3.2 (R Core Team, 2024).

#### 2.3.1 Sources of variation in temperature stress tolerance

To assess the alternative sources of temperature stress tolerance in *L. minor*, we evaluated how RGR varied between clonal lines, across temperatures and over time. To do so, we fitted a linear mixed model with restricted maximum likelihood (REML), with RGR as the dependent variable and the main effects, two-way and three-way interactions of “Clonal line”, “Temperature” and “Week” as fixed factors. Because three plants from a particular replicate were used to generate the replicate used in the following week (i.e., observations from different weeks were not independent), the identity of each replicate lineage nested within clonal line and temperature was included in the model as random factor. This allowed us to account for repeated measures across weeks without assuming the sphericity required in a repeated-measures ANOVA (Armstrong, 2017; Powers & Kozak, 2019). We accounted for any differences in light intensity associated with spatial position (i.e., proximity to the side walls where the light sources are located) by including positional grid column as a random factor.

The significance of fixed factors was evaluated using function *Anova* (package car; Fox et al., 2012) with type III sum of squares and using the Kenward–Roger approximation. Marginal and conditional *R^2^* (i.e., the variance explained only by fixed factors, and by all factors in the model, respectively), were assessed using function *r.squaredGLMM* (package MuMIn; Barton, 2020). A significant effect of “Clonal line” indicated overall genetically based differences in RGR between lines. A significant effect of “Temperature” indicated overall differences in RGR between temperature treatments (i.e., overall phenotypic plasticity to temperature). A significant effect of “Week” indicated overall variation in RGR across the duration of the experiment, reflecting changes in mean performance over time (for instance, due to either beneficial acclimation or decline under stress). A significant “Clonal line-by-Temperature” interaction indicated that clonal lines responded differently to a given temperature (i.e., G×E). A significant “Clonal line-by-Week” interaction indicated that patterns of RGR change over time differed among clonal lines. A significant “Temperature-by-Week” interaction indicated that temporal patterns of RGR change varied among temperature treatments. Finally, a significant “Clonal line-by-Temperature-by-Week” interaction indicated that clonal lines showed different patterns of temporal change in response to temperature (i.e., temporal variation in G×E). Because the “Temperature-by-Week” interaction was significant, we robustly evaluated whether RGR varied across weeks in each experimental temperature by performing a Tukey’s *post hoc* test on marginal means using the *emmeans* function (package emmeans; Lenth, 2021), which allowed to control for multiple testing within each temperature treatment.

#### 2.3.2 Associations between the performance of clonal lines and their temperature of origin

To determine whether local climatic conditions have selectively shaped the genetic variation for temperature stress tolerance in *L. minor,* we tested whether genetic variation for RGR was associated with growth-season temperature at the clone collection sites. First, we calculated the mean RGR of each clonal line in each week and temperature. We then extracted the mean temperature of the warmest quarter (bio10) in each collection site from WorldClim bioclimatic layers (Fick & Hijmans, 2017) using ArcMap 10.5 (ArcGIS Desktop, ESRI, CA, USA), employing a 2 km buffer zone around each sampling location to account for potential within-site climatic heterogeneity and coordinate imprecision. Temperature of the warmest quarter represents the local temperature during peak growth of *L. minor* in the season (i.e., in late-spring and summer). Finally, we used pairwise Pearson correlations to test whether variation in performance (RGR) within a particular week and temperature treatment was associated with variation in local temperature of origin based on the temperature of the warmest quarter; the same results were obtained using annual mean temperature (bio1; results not shown). A significant association indicated the presence of clinal variation in RGR along climatic gradients, suggesting past adaptive evolution to temperature conditions. To correct for multiple testing, significance of p-values from Pearson correlations was determined using an FDR threshold of 0.05 (Benjamini & Hochberg, 1995).

## 3. Results

We found significant main effects of clonal line, temperature, and week on relative growth rate (RGR), and all two- and three-way interactions among those factors were also significant (Table 2). Marginal *R*^2^ was extremely high (0.96), indicating that nearly all variance in RGR was explained by these factors and their interactions. Since conditional *R*^2^ was similarly high (0.98; Table 2), the random effects included in the model explained little additional variance, indicating that both replicates of each clonal line generally performed similarly and experimental position within the incubators had a minimal influence on RGR.

**Table 2:**
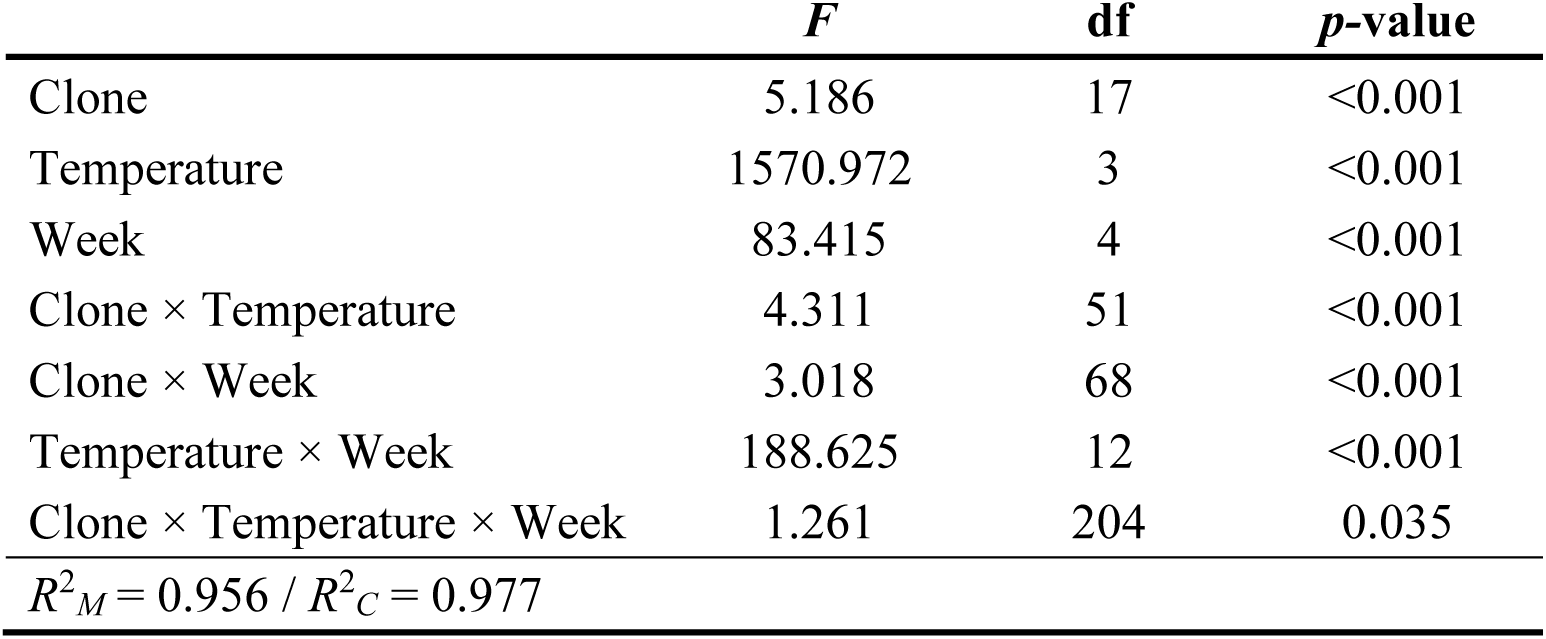
Results from the linear mixed model testing the effect of clonal line, temperature, week, and their two-way and three-way interactions on relative growth rate (RGR). Replicate position and identity (nested within clonal line) were included in the model as random factors. *F*-statistics, degrees of freedom (df) and *p-*values for each term are shown along with marginal and conditional variance of the model (*R*^2^*_M_* and *R*^2^*_C_*, respectively).

|  | *F* | df | *p*-value |
| --- | --- | --- | --- |
| Clone | 5.186 | 17 | <0.001 |
| Temperature | 1570.972 | 3 | <0.001 |
| Week | 83.415 | 4 | <0.001 |
| Clone × Temperature | 4.311 | 51 | <0.001 |
| Clone × Week | 3.018 | 68 | <0.001 |
| Temperature × Week | 188.625 | 12 | <0.001 |
| Clone × Temperature × Week | 1.261 | 204 | 0.035 |
| $R^2_M = 0.956 / R^2_C = 0.977$ | | | |

Our results showed overall RGR responses to temperature (significant main effect of “Temperature”; Table 2). Relative growth rate was highest and similar at 23 and 28 °C (grand mean of 0.309 cm^2^/day), declining by 28% at 18°C (0.223 cm^2^/day) and 85% at 33 °C (0.045 cm^2^/day). As noted above, we also detected average genetic differences among clonal lines in RGR (significant main effect of “Clonal line”; Table 2). However, the significant interactions of “Clonal line” with “Temperature” and “Week”, along with the three-way interaction, indicated that genetic differences in performance varied with temperature and over time (Table 2; Fig. 2).

**Figure 2:**
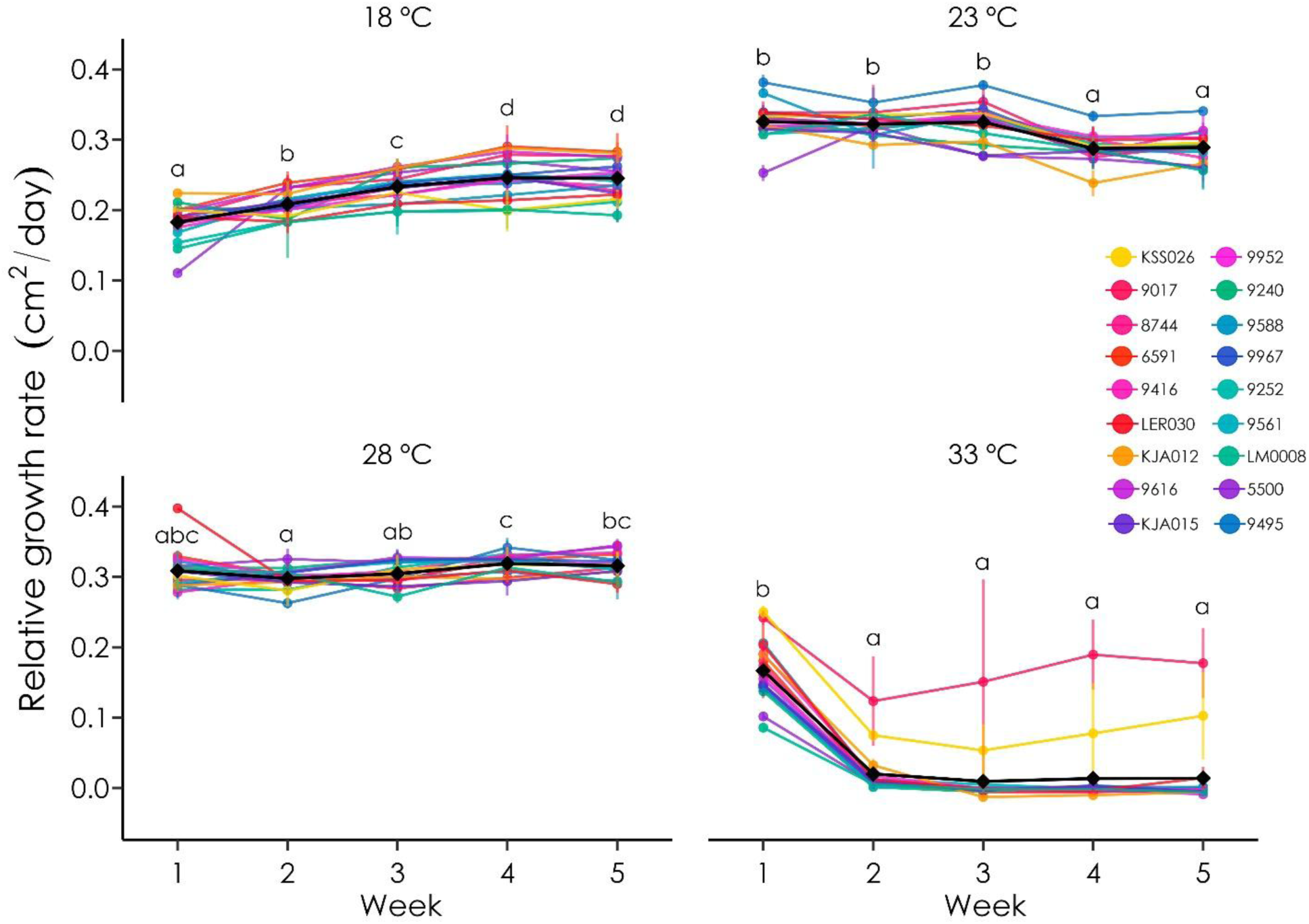
Relative growth rate (RGR) variation among clonal lines of *Lemna minor* over time (week 1 – week 5) within each temperature treatment. Each colored line represents the temporal trajectory of a single (color-labeled) clonal line at a specified temperature. Black lines and diamonds represent the mean temporal trajectory and overall mean performance of clonal lines within each temperature. Different letters indicate significant differences between weeks in a particular temperature, based on Tukey-adjusted post hoc tests (see *Methods)*.

Overall RGR (i.e., averaged across clones) varied over time (main effect of “Week”; Table 2), but in a temperature-dependent manner (“Temperature-by-Week” interaction; Table 2). Specifically, average RGR increased significantly over time (an indicator of physiological acclimation) only at 18 °C (35% increase from week 1 to week 5; Fig. 2). In contrast, at 23 °C post-hoc tests showed an 11% average reduction in RGR between the first three and the last two weeks of the experiment, and at 28 °C we found no significant differences in RGR between weeks 1 and 5 (Fig. 2). Finally, at 33 °C RGR dropped sharply during the experiment, with a significant, drastic average reduction of 88% after the first week.

After a week in this heat-stress treatment, most clonal lines exhibited virtually no growth, yet those from the two warmest sites (KSS026 and 9017) expressed a distinctive response, maintaining relatively high RGR values throughout the 5-week experiment (significant “Clonal line-by-Temperature-by-Week” interaction; Table 2, Figs. 2-4). Importantly, while there was generally very high congruence between replicate observations in our experiment (see above), at 33° C replicates within these two lines differed markedly, with one of the two replicates showing a sharp RGR reduction and subsequent partial recovery, and the other replicate maintaining relatively high growth throughout the experiment (Fig. 3).

**Figure 3:**
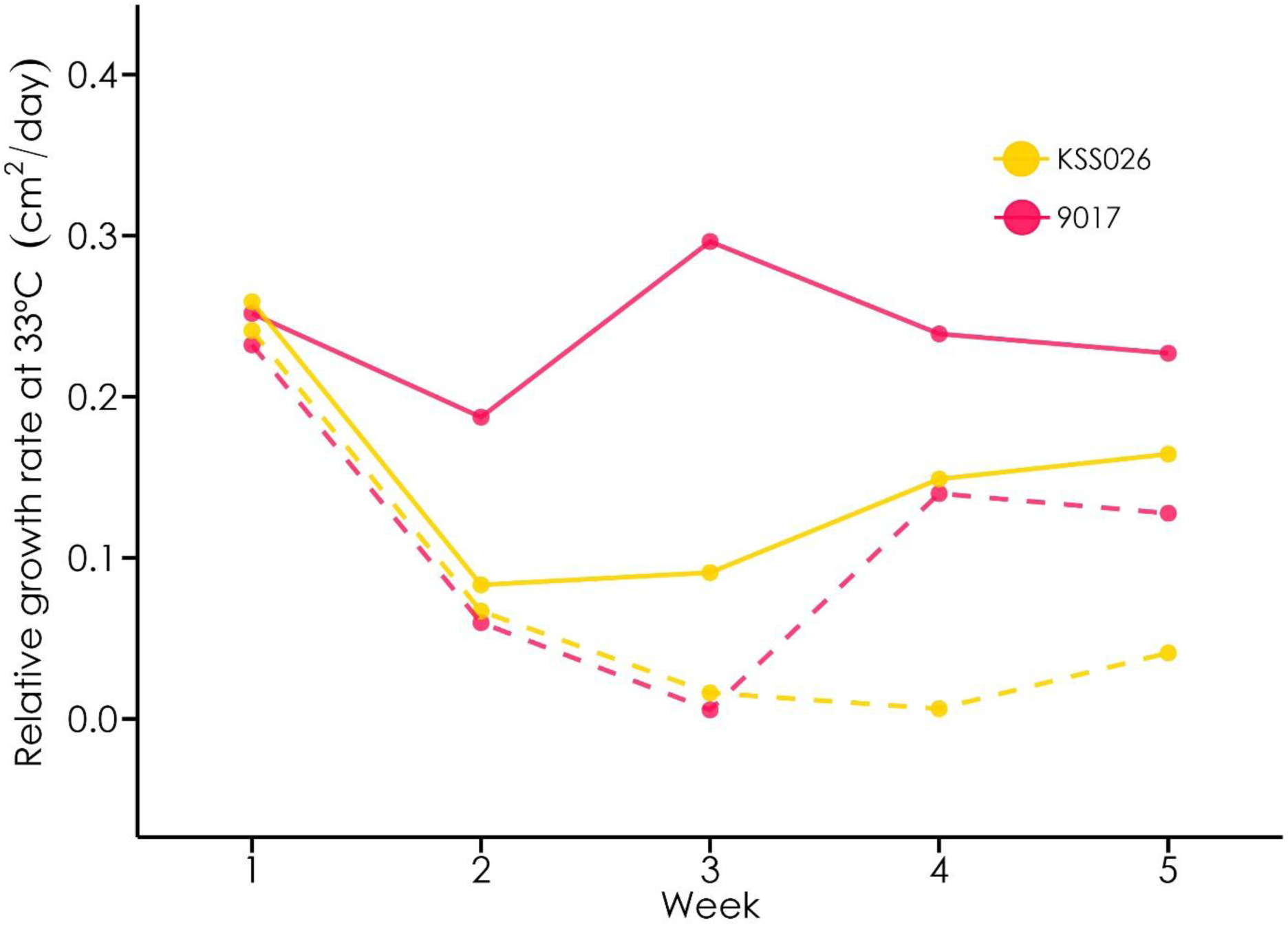
Temporal variation in relative growth rate (RGR) at 33 °C for the two most heat-tolerant clonal lines (KSS026 and 9017). Solid and dashed lines represent the two experimental replicates of each clonal line. Divergent trajectories between replicates within the same clonal line show within-line temporal variation in performance under prolonged heat exposure.

**Figure 4:**
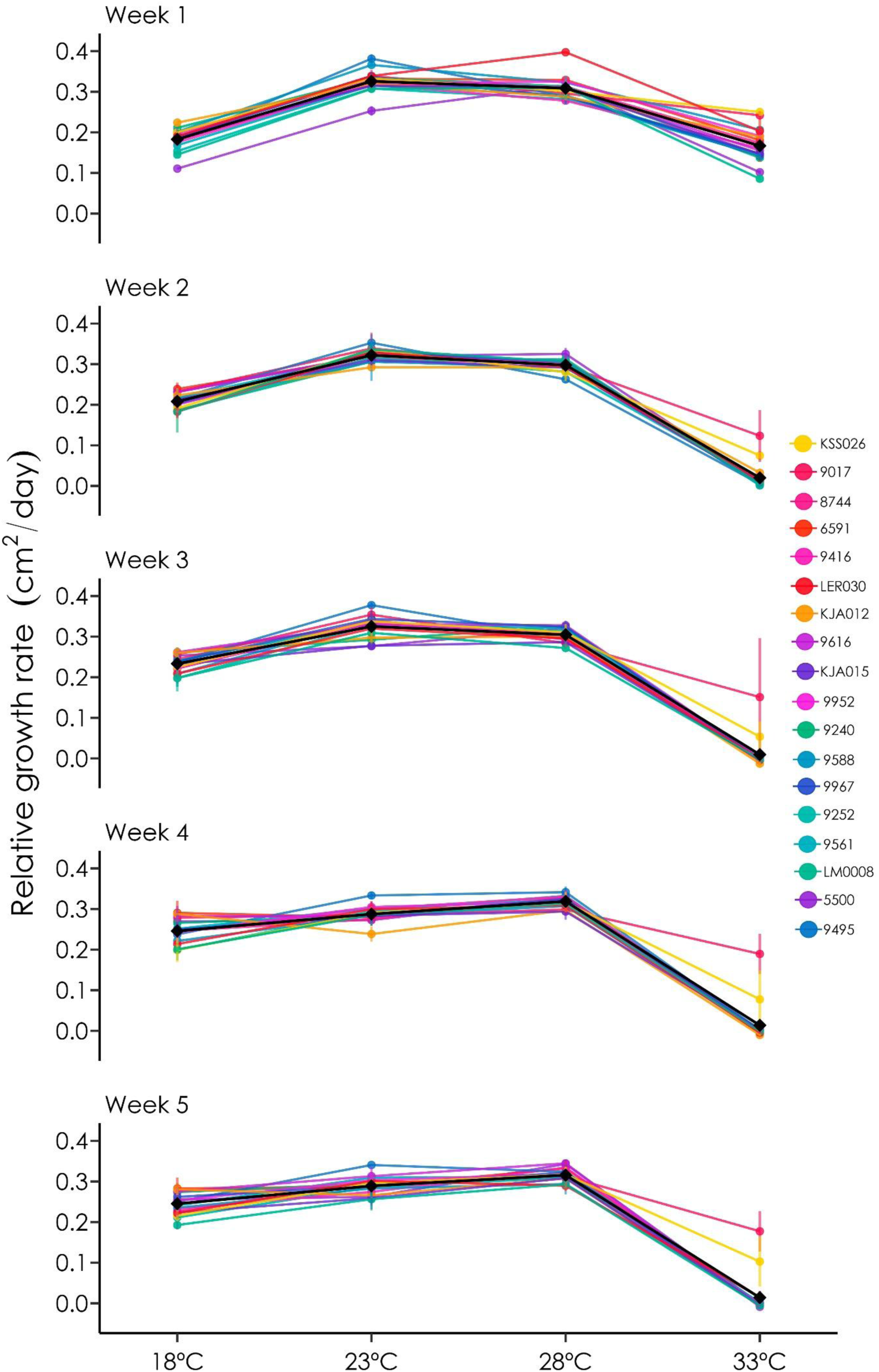
Variation in relative growth rate (RGR) of clonal lines across temperatures within each experimental week. Each colored line represents the temperature response of a clonal line at a given week. Black lines and diamonds represent overall performance and temperature response of all clonal lines within each week.

In view of the significant three-way interaction between “Clonal Line”, “Temperature”, and “Week”, we evaluated possible adaptive differentiation between lines by performing Pearson correlations between RGR in each week and experimental temperature, and the local climate (temperature of the warmest quarter) of the experimental lines. We detected a significant RGR-climate correlation at 33°C but only in the first week, when clonal lines from warmer sites showed significantly higher RGR after FDR correction (Fig. 5). Although this pattern suggests that clonal lines from warmer climates were generally more tolerant of heat stress, this advantage disappeared after the first week.

**Figure 5:**
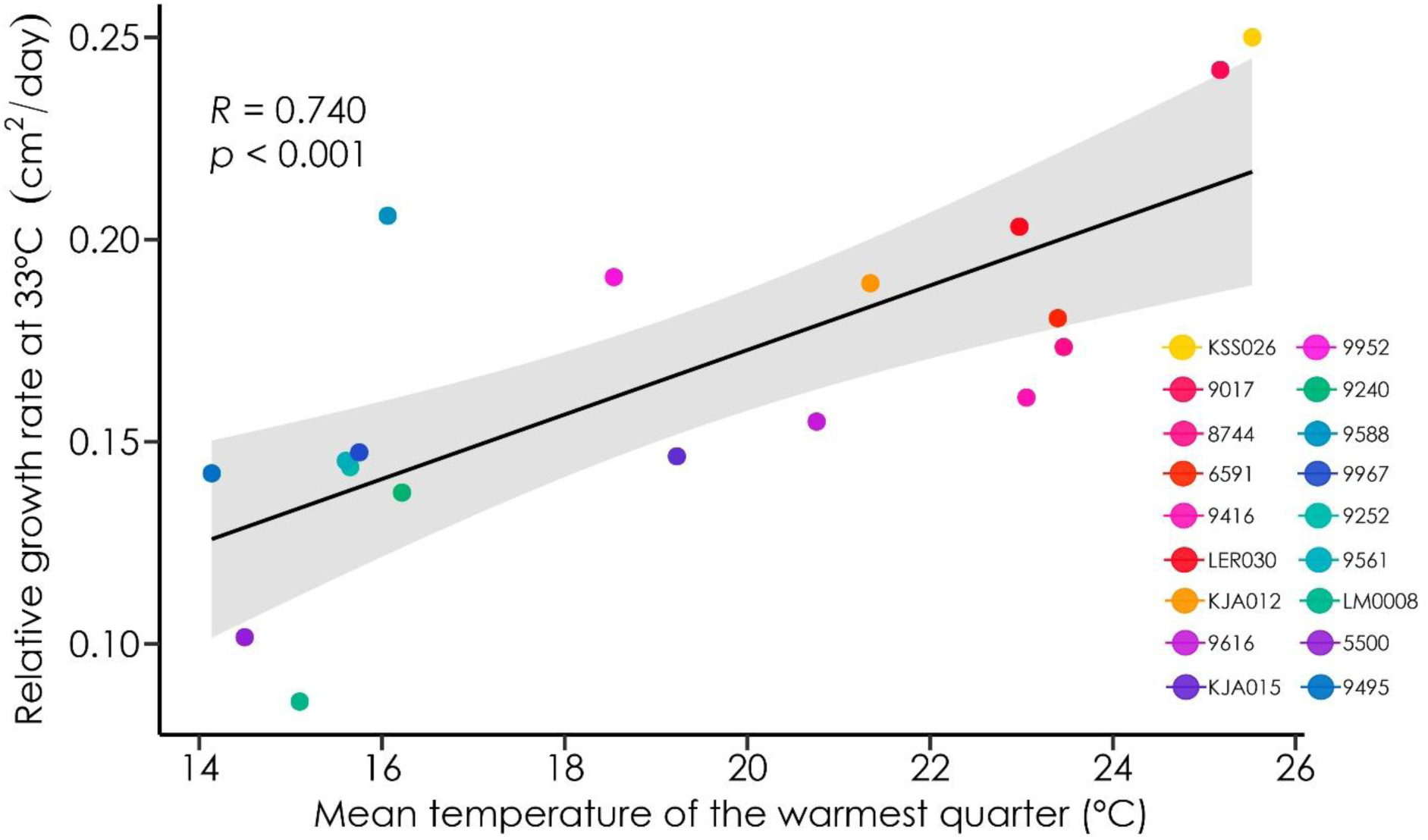
Association between mean temperature of the warmest quarter at the collection site and performance (RGR) at 33 °C of the clonal lines, in week 1. The regression line represents the Pearson correlation, with shaded areas indicating 95% confidence intervals. Pearson correlation coefficient (*R*) and *p*-value after FDR correction are shown. Due to minimal variation between the two replicates of the same line, standard errors are not visible.

## 4. Discussion

Our study evaluated adaptive variation for temperature stress tolerance in *L. minor* to shed light on its drivers and potential eco-evolutionary consequences in the context of climate change. The results indicate that variation in heat stress tolerance reflects several non-exclusive mechanisms that operate at different ecological and temporal scales. First, contrary to previous evidence in duckweeds, we detected adaptive genetic differentiation in heat stress tolerance among clonal lines related to their climate of origin. However, these adaptive differences among lines were expressed only transiently, diminishing under prolonged exposure to heat stress. Thus, genetic differences in *L. minor* tolerance to heat stress are not fixed but rather reflect evolved differences in the expression of temperature- and duration-dependent responses. Additionally, we identified two further dynamic aspects of *Lemna* growth response to temperature: (i) physiological acclimation to certain temperatures; and (ii) within-line variation expressed under sustained heat stress.

The common freshwater species *L. minor* encounters wide climatic variation across its global distribution, which likely imposed differential selective pressures among its populations (Ziegler *et al*., 2015). Our findings provide the first evidence of adaptive genetic differentiation in heat stress tolerance associated with climatic variation in this species. Clonal lines from warmer sites initially showed significantly higher performance under heat stress than lines from colder regions (Fig. 5). These results suggest that populations of *L. minor* have undergone past adaptation to their local temperatures, consistent with previous evidence for divergent evolution in response to climatic variation in other aquatic and terrestrial plant species (e.g., Gillard *et al*., 2017; Solé-Medina *et al*., 2022; Blanco-Sánchez *et al*., 2024). Although such genetically-based differences in heat stress tolerance may confer a higher ability to persist under climate change conditions to the lines from warmer regions, we note that the differences were expressed only during the initial phase of stress exposure. Sustained heat stress greatly reduced adaptive differences among clonal lines, and even most of the more tolerant lines could not maintain their RGR throughout the five-week experiment. This pattern is consistent with the idea that organisms in warmer environments operate close to their thermal tolerance thresholds, such that even small increases in temperature can result in sharp performance declines, potentially leading to similar or even greater vulnerability (Somero, 2010; Bennett *et al*., 2019; Sage, 2020). Nevertheless, the higher ability to withstand acute temperatures for 1-2 weeks that we observed in clones from warm climates, even when it does not extend to longer periods, could provide substantial advantages for coping with relatively long heatwaves.

Interestingly, previous studies have failed to detect clinal variation in *L. minor*, attributing these results to the predominant role of neutral evolutionary processes such as migration and colonization events in shaping genetic variation (Honnay *et al*., 2010; Senevirathna *et al*., 2023). These results have been attributed to the clonal reproduction of duckweed species and the complex demographic processes experienced by aquatic plants. Because seeds and pollen are rarely produced under natural conditions, colonization and genetic admixture of duckweed populations rely on the endo- and epizoochorous dispersion of duckweed fronds by waterfowl, their main dispersal vectors (Coughlan *et al*., 2015; Silva *et al*., 2018; Tippery & Les, 2020; Paolacci *et al*., 2023). Historical demographic processes due to limited and uneven dispersal often result in strong founder effects and enhanced likelihood of genetic drift, and these non-adaptive evolutionary processes can significantly shape genetic (and hence phenotypic) variation within and among populations (Keller & Taylor, 2008; Sork, 2016; Ramos-Muñoz *et al*., 2024). Consequently, clonal species generally show limited within-population genetic variation, which in turn significantly constrains adaptation via natural selection (Dodd & Douhovnikoff, 2016; Sammarco *et al*., 2022). Indeed, low levels of standing genetic variation have been previously reported in duckweed populations (Jordan *et al*., 1996; Xu *et al*., 2019; Gillies *et al*., 2024). Nevertheless, our findings (and those in some other predominantly asexual species; Woudstra *et al*., 2024) indicate that adaptive differentiation can still emerge when clonal lineages have evolved in consistent, contrasting environmental conditions.

Our results further suggest that previous failures to detect adaptive differentiation in *L. minor* may reflect context-dependent expression of genetic differences rather than a true absence of climate-related differentiation. In our experiment, a significant association between RGR and the climate of origin was detected only under a specific combination of stress intensity and exposure time (i.e., only in the first week and at 33°C; Fig. 5). It is well known that the expression of quantitative genetic variation depends on environmental conditions (see Hoffmann & Merilä, 1999; Charmantier & Garant, 2005; Ghalambor *et al*., 2007; Sultan, 2007; van Kleunen & Fischer, 2008), and indeed, many previous studies have identified genetic differentiation only under particular conditions (e.g. Ramírez-Valiente *et al*., 2018, 2022; Cooper *et al*., 2022). Our results extend this point by showing that the detectability of adaptive differentiation can also depend on the duration of exposure. In most studies with duckweeds, plants are phenotyped after long-term exposures to constant conditions (e.g., Van Antro *et al*., 2023; Couture *et al*., 2025), which may obscure footprints of selection when adaptive differences are expressed only transiently or under specific environmental conditions. Our findings emphasize that detecting adaptive phenotypic variation requires its evaluation under ecologically and temporally relevant conditions.

Similar to patterns of adaptive differentiation, acclimation was also temperature-dependent in *L. minor*. Acclimation (i.e., physiological adjustments that enhance performance over time within a given environment; IUPS Thermal Commission, 2001; Bowler, 2005; Loeschcke & Sørensen, 2005) was observed at 18 °C, but not at temperatures closer to the species’ optimum (23 and 28 °C). In plants, thermal acclimation can include several responses such as regulatory changes in gene expression, induction of heat-shock proteins, stabilization of cellular membranes, and adjustments in photosynthesis and respiration rates, among others (Atkin & Tjoelker, 2003; Way & Yamori, 2014; Balfagón *et al*., 2020). Although we cannot pinpoint the exact physiological responses behind such differences in acclimation across temperatures, it is worth noting that clonal lines were pre-acclimated to 22 °C before the experiment. Thermal acclimation benefits are not expected to be ubiquitous but context-specific (Smith *et al*., 2021), increasing over time under conditions that clearly differ from those experienced recently in the development (Semsar-kazerouni & Verberk, 2018). Therefore, beneficial acclimation would have occurred during such pre-acclimation phase, making them undetectable in our experiment at temperatures close to the pre-experiment culturing conditions such as 23 °C. Furthermore, since 23-28 °C matches the optimum range of temperatures for *L. minor* (Lasfar *et al*., 2007), long-term selection may have previously canalized physiological performance, reducing both the ability and the need for further acclimatory adjustments to these conditions (Armitage & Jones, 2019; Van Dyck *et al*., 2021). In contrast, 18 °C corresponds to the low end for the growth of the species, especially for lines sensitive to cold stress. These results suggest that acclimation may arise more frequently under sub-optimal yet not highly stressful conditions, consistent with the view that physiological adjustments are expected to be more costly (and therefore selected against) under higher stress (Valladares *et al*., 2007; Stotz *et al*., 2021; Solé-Medina *et al*., 2022; but see Ramos-Muñoz *et al*., 2025).

Our results also suggested a third, unexpected dimension of adaptive variation in thermal stress tolerance in *L. minor*, potentially related to epigenetic mechanisms: functional differences in heat stress tolerance patterns between replicates of the same clonal line. As noted above, while most of the clonal lines collapsed at 33°C between the second and the third week of the experiment, the two most heat-tolerant lines (KSS026 and 9017, collected from the warmest sites) maintained moderate to high RGR across all five weeks (Figs. 2 and 3). Yet, in contrast to the consistent response trajectories observed between replicates of the other clonal lines, the two replicates of each of these lines differed notably in their performance over time. In both cases, one replicate from each line showed constant positive RGR, while the other suffered a steep initial RGR decrease followed by a partial recovery later (Fig. 3; see also error bars in Fig. 2). Such pronounced variation in environmental response between replicates of the same line suggests the existence of an independent acclimatory process that can differ between genetically identical individuals. One potential process involved could be epigenetic marks that may be maintained stably across clonal generations (Richards *et al*., 2017; Shahmohamadloo *et al*., 2025) via such mechanisms as DNA methylation, histone modifications and the transmission of small RNAs (reviewed in Adrian-Kalchhauser *et al*., 2020). Among these mechanisms, DNA methylation includes stochastic and environmentally induced components (Sammarco *et al*., 2024; Shahmohamadloo *et al*., 2025), which can cause heritable and functional variation between isogenic replicates (Verhoeven *et al*., 2010; Lemmen *et al*., 2022; Sammarco *et al*., 2022; D’Aguillo & Sultan *in prep.*). Indeed, stressful conditions often increase the rate of epimutations (Verhoeven & Preite, 2014), and exposure to high temperature was previously shown to induce heritable DNA methylation variants in *L. minor* (van Antro et al. 2023). Such epigenetic variability might function as a form of bet-hedging that increases the likelihood of persistence of some individuals under unpredictable or stressful conditions (Herman *et al*., 2014; Shahmohamadloo *et al*., 2025). Thus, genetic differences in epigenetic capacity (Adrian-Kalchhauser *et al*., 2020), together with environmentally and stochastically-induced epigenetic marks, may jointly contribute to variation in heat stress tolerance in *L. minor*.

## 5. Conclusions

Our findings reveal that variation in temperature stress tolerance in *L. minor* emerged from multiple interacting sources, rather than a single underlying process. Historical exposure to contrasting temperature conditions led to adaptive genetic differentiation among clonal lines, shaping initial variation in heat stress tolerance. Short-term environmental conditions also modulated performance through temperature-dependent acclimation. Finally, novel patterns of adaptive variation emerged under sustained heat stress within certain clonal lines, potentially mediated by epigenetic mechanisms. Together, our results highlight that variation in heat stress tolerance is not a fixed property of genotypes, but a dynamic outcome arising from the interplay between past selection, immediate physiological adjustments, and newly generated phenotypic variation under stress. This multi-layered view provides a more nuanced framework for understanding plant responses to increasing temperature stress under ongoing climate change.

## Acknowledgements

We thank Nele Horemans (SCK CEN; Belgium), Manuela Bog and Klaus Appenroth (Friedrich Schiller University Jena; Germany), and Laura Morello, Silvia Giani and Luca Braglia (IBBA CNR; Italy) for providing us with the clonal lines used in this study and their suggestions about how to culture the plants. We are also indebted to Meret Huber and Shuqing Xu for the insightful discussions. Finally, special thanks to Slavica Milanovic-Ivanovic and Gregor Disveld for their technical support during the experiment.

## CRediT authorship contribution statement

**Mario Blanco-Sánchez:** Conceptualization, Methodology, Formal Analysis, Investigation, Data Curation, Writing - Original Draft, Visualization. **Sonia E. Sultan:** Conceptualization, Writing - Review & Editing. **Koen J. Verhoeven:** Conceptualization, Methodology, Writing - Review & Editing, Supervision, Project administration, Funding acquisition.

## Data accessibility statement

The data generated in this study will be uploaded to a public repository upon manuscript acceptance.

